# Artificial Gravity During a Spaceflight Analog Alters Brain Sensory Connectivity

**DOI:** 10.1101/2022.11.02.514887

**Authors:** Heather R. McGregor, Jessica K. Lee, Edwin R. Mulder, Yiri E. De Dios, Nichole E. Beltran, Scott J Wood, Jacob J. Bloomberg, Ajitkumar P. Mulavara, Rachael D. Seidler

**Author notes:** **Corresponding author:** Rachael D. Seidler, Ph.D., University of Florida, 1864 Stadium Rd., Gainesville, FL 32611. **Author Contributions:** HRM analyzed data. JKL collected data. JKL, HRM, YED, NEB, ERM, RDS set up the experiment. RDS, ERM, JJB, APM, SJW designed the experiment and secured funding. HRM drafted the manuscript. HRM, JKL, ERM, JJB, RDS edited the manuscript. **Role of the Funder/Sponsor:** The funding sources had no role in the design and conduct of the study; collection, management, analysis, and interpretation of the data; preparation, review, or approval of the manuscript; and decision to submit the manuscript for publication. **Data Availability Statement:** MRI files for this study will be placed in the NASA data repository upon study completion.

## Abstract

Spaceflight has numerous untoward effects on human physiology. Various countermeasures are under investigation including artificial gravity (AG). Here, we investigated whether AG alters resting-state brain functional connectivity changes during head-down tilt bed rest (HDBR), a spaceflight analog. Participants underwent 60 days of HDBR. Two groups received daily AG administered either continuously (cAG) or intermittently (iAG). A control group received no AG. We assessed resting-state functional connectivity before, during, and after HDBR. We also measured balance and mobility changes from pre-to post-HDBR. We examined how functional connectivity changes throughout HDBR and whether AG is associated with differential effects. We found differential connectivity changes by group between posterior parietal cortex and multiple somatosensory regions. The control group exhibited increased functional connectivity between these regions throughout HDBR whereas the cAG group showed decreased functional connectivity. This finding suggests that AG alters somatosensory reweighting during HDBR. We also observed brain-behavioral correlations that differed significantly by group. Control group participants who showed increased connectivity between the putamen and somatosensory cortex exhibited greater mobility declines post-HDBR. For the cAG group, increased connectivity between these regions was associated with little to no mobility declines post-HDBR. This suggests that when somatosensory stimulation is provided via AG, functional connectivity increases between the putamen and somatosensory cortex are compensatory in nature, resulting in reduced mobility declines. Given these findings, AG may be an effective countermeasure for the reduced somatosensory stimulation that occurs in both microgravity and HDBR.

## INTRODUTION

Human spaceflight is associated with many physiological changes, including effects on both the nervous system and behavior. For instance, balance and mobility declines are evident immediately post-flight and last for several days to weeks as crewmembers readapt to Earth’s gravity ^1–3^. These changes are thought to reflect aftereffects of adaptation to the altered sensory signaling that occurs in microgravity, including altered vestibular inputs due to the lack of gravitational forces acting on the otoliths and reduced somatosensory signaling due to body unloading. Studies have reported changes in brain structure ^4–10^ and function ^11–17^ inflight and post-flight as well. Some of these changes seem to reflect the physical effects of the microgravity environment, including headward fluid shifts and an upward shift of the brain within the skull ^4,5^. Other brain changes with spaceflight have been suggested to reflect neuroplasticity and functional adaptations to the microgravity environment; this interpretation is supported by correlations between these brain and behavior changes occurring inflight or post-flight ^9,11,18^.

Several studies have used long duration (days to months) exposure to head down tilt bed rest (HDBR) to mimic some of the effects of microgravity. Indeed, this environment results in similar effects to microgravity such as reduced sensory inputs to the bottom of the feet, axial body unloading, and headward fluid shifts ^19,20^. HDBR is also associated with subsequent postural and mobility declines as well as altered brain activity ^2,20–26^.

Resting state functional MRI — an approach for estimating correlated neural activity across distributed brain networks ^27^ — has revealed changes in brain network functional connectivity (FC) with HDBR ^24,25,28–32^. Studies of short (72 hours ^22,30^) and longer (30-70 days ^24–26,31,32^) duration HDBR have revealed changes in FC from pre-to post exposure. For example, we previously found that 70 days of HDBR resulted in increased FC between motor and somatosensory brain regions; participants who showed FC increases between these regions the most had the smallest decrements in balance after exiting HDBR ^26^. These FC changes may reflect sensory reweighting that occurred during HDBR which was subsequently beneficial for balance. Interestingly, a single subject case study of a cosmonaut reported decreased resting state connectivity within the right insula (overlapping with vestibular cortex) from pre-to post-spaceflight ^12^. We also found increased connectivity between the vestibular cortex (retroinsular cortex and parietal opercular area 2, OP2) and the cerebellum with HDBR, and decreased connectivity between left and right temporoparietal junctions (overlapping with vestibular cortex ^26^). While it may seem unusual that vestibular cortical connectivity changes in HDBR even though there are no gravitational changes, there is a rotation of the body’s typical orientation relative to the gravitational vector, and there is also evidence for multisensory reweighting in this environment ^20^. We found similar results in another bed rest study with elevated CO^2^ (to mimic levels seen in the enclosed environment of the International Space Station ^24^).

Numerous potential countermeasures have been investigated to mitigate the physiological changes that occur with human spaceflight, including exercise ^2,33^ and lower body negative pressure ^34,35^. Exercise performed aboard the ISS does not completely prevent spaceflight-induced sensorimotor changes, however ^2^. One countermeasure approach that is appealing due to its broad, systemic impact is artificial gravity (AG) ^36^; it engages muscles, loads bones and the cardiovascular system, and stimulates the otoliths to what occurs due to Earth’s gravity. AG can be achieved in space with either a rotating craft or a short-arm centrifuge. HDBR studies have also investigated the effects of AG by having participants undergo daily centrifugation as a potential countermeasure. A 21-day HDBR investigation reported that participants that received daily AG shifted their position perception to be 30° closer to upright than those that did not ^37^. A more recent study found that intermittent (six bouts of five minutes each) AG reduced HDBR-associated orthostatic intolerance following five days of HDBR more than a continuous daily session of 30 minutes ^38^. Behavioral changes during the current bed rest campaign have been recently reported ^39,40^, showing that 30 minutes of daily AG offered modest or no mitigation ^39,40^ of mobility and balance declines that typically accompany HDBR ^1,41,42^. Though there were group differences in motion sickness throughout bed rest, with subjects in the continuous AG group reporting more severe motion sickness symptoms than both the intermittent AG group and the non-centrifuged control group ^40^. Despite modest or no group differences in performance changes following bed rest, it is unknown whether the groups showed differences in functional brain changes with HDBR, perhaps reflecting compensation.

In the current paper, we report the effects of 30 minutes of daily AG applied along the long axis of the body, with 1 g at the center of mass, during a 60 day HDBR study. We investigated changes in resting-state brain functional connectivity occurring with HDBR and whether they differed between participants that did versus did not receive daily AG. We also tested for alterations in brain-behavior correlations. We hypothesized that the vestibular and somatosensory stimulation that occur with AG would alter HDBR-associated changes in sensorimotor connectivity. We also expected that these changes would correlate with mobility and balance changes occurring from pre-to post HDBR.

## METHODS

### Participants

This study was carried out as part of the AGBRESA (Artificial Gravity Bed Rest study with European Space Agency) bed rest campaign in 2019. It was conducted at the German Aerospace Center’s :envihab facility in Köln, Germany.

Prior to enrolment, prospective volunteers were prescreened for being able to tolerate AG, which was applied here using a short-arm centrifuge such that participants received 1 g at their center of mass, directed axially towards the feet. This resulted in a centrifuge speed of approximately 30 revolutions per minute, depending on participant height. Twenty-four individuals participated after providing their consent (8 F, age: 33.3 ± 9.17 years, height: 174.6 ± 8.6 cm, weight: 74.2 ± 10.0 kg). The study was approved by the NASA and University of Florida Institutional Review Boards, as well as the local ethical commission of the regional medical association (Ärztekammer Nordrhein). Three participants withdrew early in the study and were replaced; we do not present their partial datasets here.

### Experimental Design

Participants resided at the :envihab facility throughout this 88 day study (Figure 1A). All participants underwent an ambulatory 14-day baseline data collection (BDC) phase prior to commencing a 60 o-day phase of strict 6° head down tilt bed rest (HDBR). Participants remained in the head down tilt position at all times during the HDBR phase (all activities and hygiene maintenance took place in this position) and were monitored for compliance. On the first day of HDBR, participants were assigned to one of three groups: i) the continuous AG (cAG, n=8) group exposed to a continuous 30-minute bout of centrifugation each day of HDBR, ii) the intermittent AG group (iAG, n=8) exposed to six ⍰05-minute bouts of centrifugation each day of HDBR, or iii) the control group (CTRL, n=8) that received no centrifugation during HDBR. AG was applied using a short-arm centrifuge such that participants received 1 G at their center of mass, directed axially towards the feet. This resulted in a centrifuge speed of approximately 30 revolutions per minute, depending on participant height. After the 60-day bed rest phase, participants underwent a 14-day recovery phase during which they were re-ambulated within the bed rest facility.

**Figure 1.**
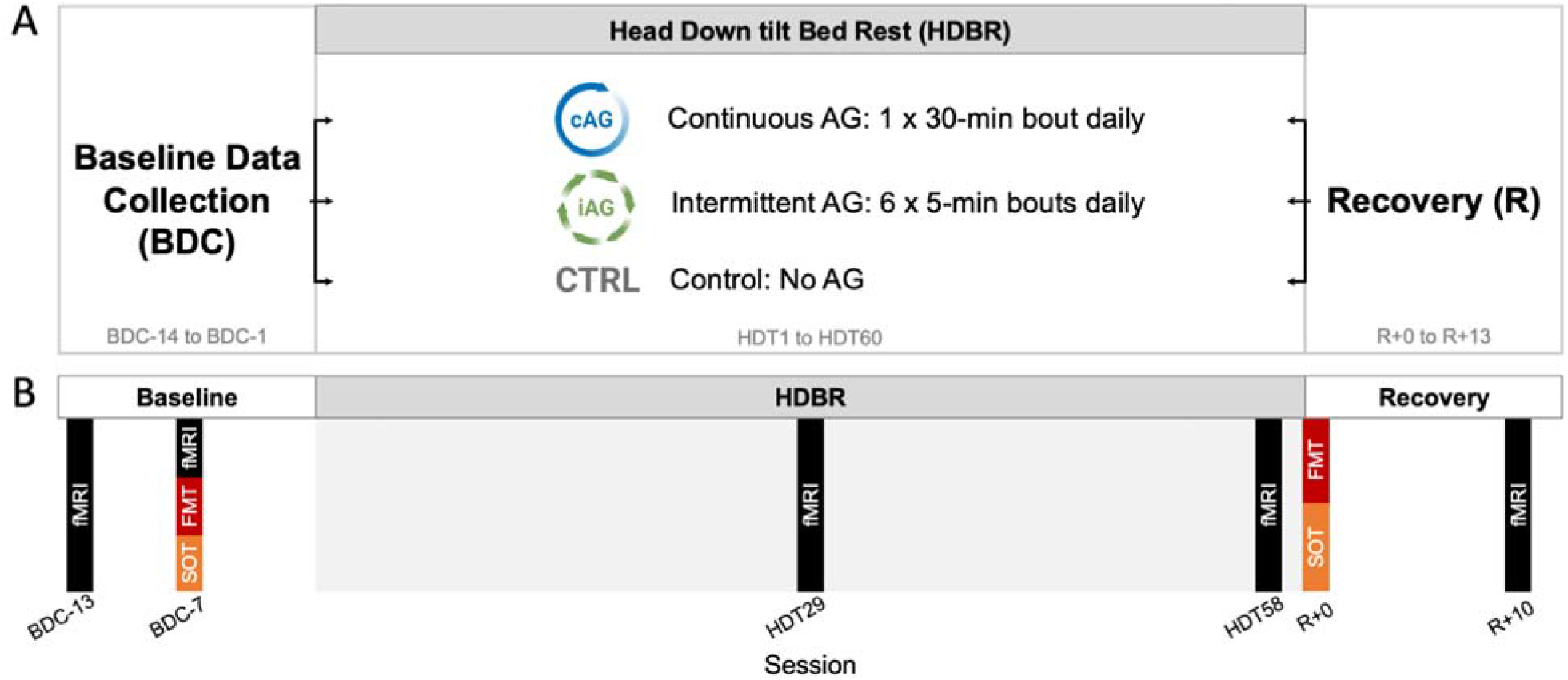
Experimental Design. **A)** Baseline data collection (BDC) occurred during a 14-day ambulatory phase. All participants then underwent 60 days of strict 6° head down tilt bed rest (HDBR), assigned to either the continuous AG (cAG), intermittent AG (iAG), or control (CTRL) group. Following the bed rest intervention phase, all participants underwent a 14-day ambulatory recovery (R) phase. **B**) Resting-state fMRI scans were acquired twice during the BDC phase (BDC-13, BDC-7), twice during the HDBR phase (HDT29, HDT58), and once during the recovery phase (R+10). Mobility and balance were assessed using Functional Mobility Tests (FMT) and Sensory Organization Tests (SOTs). These behavioral tests were performed on BDC-7 and after exiting bed rest on R+0. Familiarization of behavioral tests occurred on BDC-13 (not shown). AG, artificial gravity.

### MRI Data Collection

As shown in Figure 1B, MRI data were collected during 5 scan sessions: twice during the BDC phase 13 and 7 days prior to bed rest (i.e., BDC-13 and BDC-7, respectively), on days 29 and 58 of HDBR (i.e., HDT29 and HDT58, respectively), and once during the recovery phase 10 days after the end of bed rest (i.e., R+10).

All MRI scans were collected using a 3T Siemens Biograph mMR scanner. While in the scanner, participants laid on a foam wedge to maintain their 6° HDT position, but their head was positioned horizontally (0°) inside a 16-channel head/neck coil. Anatomical images were collected during each scan session with the following T1-weighted MPRAGE sequence: TR = 1900 ms, TE = 2.44 ms, flip angle = 9°, iPAT acceleration factor = 2, FOV = 250 × 250 mm, matrix = 512 × 512, voxel size = 05. x 0.5 × 1.00 mm with 0.5 mm slice gap, number of slices = 192. Resting-state fMRI data were collected with a T2*-weighted echo planar imaging sequence with TR = 2500 ms, TE = 32 ms, flip angle = 90°, FOV = 192 × 192 mm, matrix = 64 × 64, voxel size 3 × 3 × 3.5 mm, number of slices = 37, duration = 7 minutes. Participants were instructed to gaze at a red fixation circle on the screen and to let their mind wander, not thinking about anything in particular.

### Behavioral Tests

#### Functional Mobility Test

We assessed HDBR-induced changes in subjects’ whole-body locomotor function using the Functional Mobility Test (FMT). The FMT is a 6 × 4 m obstacle course ^43^. The first half was set up on a hard floor while the second half was set up upon a medium density compliant foam surface to perturb reliability of proprioceptive inputs ^43^. Starting from a seated position, the subjects must walk around, under and over a series of obstacles. Subjects completed 10 trials of the FMT by walking quickly without touching the obstacles. Subjects were familiarized with the FMT by performing 10 practice trials on BDC-13; these data were not included in the current analysis. Subjects performed the FMT on BDC-7 (pre-bed rest) and on R+0 (the day they exited bed rest) (see Figure 1B). Here we investigated HDBR-induced mobility changes by examining changes in the total completion time of the first FMT trial per session. Performance on the first FMT trial of each session is the most sensitive to post-bed rest changes in mobility as opposed to motor learning effects across FMT trials.

#### Sensory Organization Tests

We assessed changes in postural control with HDBR using Sensory Organization Tests (SOTs). SOTs were conducted using Computerized Dynamic Posturography (EquiTest, NeuroCom International, Clackamas, OR, United States) ^20^. Subjects maintained a quiet stance on a sway-referenced platform with their eyes closed. For the SOT-5 test, subjects held their head upright and stationary. For the SOT-5M test, subjects performed dynamic ±20° head pitches at a rate of 0.33 Hz paced by an auditory signal (i.e., a continuous sinusoidal wavetone). Equilibrium Quotient scores were calculated to quantify the peak-to-peak center of mass sway angle in the anteroposterior direction ^44^. Equilibrium Quotient scores range between 0 and 100 where 0 reflects an impending fall (with a 12.5° theoretical limit of stability) and 100 reflects no anteroposterior sway. Subjects performed three 20-s trials of each SOT. We used the median scores across the three trials per session to assess changes in postural stability with HDBR.

#### Image Preprocessing

We first estimated and corrected for magnetic field inhomogeneities in the resting-state fMRI scans. Prior to each resting-state run, we acquired 2 echo-planar images of opposite phase encoding direction, which we used to generate per-session field maps using topup ^45^ from the FMRIB Software Library (FSL, https://fsl.fmrib.ox.ac.uk/fsl/). Each field map was then inputted into the FieldMap Toolbox ^46^ to calculate a voxel displacement map. Subsequent preprocessing was conducted using the CONN toolbox version 18a ^47^ implemented in Matlab 2018b. Resting-state fMRI images were slice-timing corrected, realigned to the first volume, unwarped using the voxel displacement map, and resliced to 2 mm isotropic voxels. We then utilized the ARTifact detection tool (ART; http://www.nitrc.org/projects/artifact_detect) to examine volume-wise displacement and the global brain signal. We used a thresholding set at the 97th percentile settings with a within-run composite movement threshold of 0.9 mm and a global mean intensity threshold at 5 standard deviations of the mean image intensity). Volumes exceeding these thresholds were considered outliers and were included in a “scrubbing” regressor used as a covariate during denoising steps (see below).

#### Image Normalization

The brain undergoes gross morphological changes during both spaceflight ^10,48,49^ and bed rest analogs ^50^. To improve our within-subject longitudinal normalization to MNI standard space, we used a stepwise procedure using Advanced Normalization Tools (ANTs) ^51^ as in our previous spaceflight ^4,11,14^ and bed rest studies ^52,53^. First, we generated subject-specific structural templates by coregistering and averaging the subject’s 5 native space skull-stripped T1 images using antsMultivariateTemplateConstruction.sh with affine and symmetric normalization (SyN) registration. We created subject-specific functional templates using the mean of each preprocessed resting-state run. Each subject’s structural and functional templates were co-registered using rigid and affine registration. Next, the subject-specific structural template was then registered to a 1 mm-resolution MNI space template using rigid, affine, and SyN registration. We concatenated the transformations yielded by the registration steps above into a single flow field for each subject, session, and image modality. This enabled us to transform each native space preprocessed image (T1 or resting-state fMRI run) into MNI space in a single step. MNI-normalized T1 and resting-state images were then entered into CONN. Resting-state data were spatially smoothed using a 5 mm full width at half maximum (FWHM) three-dimensional Gaussian kernel.

#### Image Denoising

We estimated confounding noise arising from non-neuronal sources using the anatomical component-based noise correction (aCompCor) tool ^54^ within CONN. This method extracts signals from within eroded white matter and CSF masks applied to unsmoothed resting state fMRI data. These signals were entered into a principal components analysis, with the top five components for white matter and CSF signals reflecting noise and potential artifacts from heart beats and respiration. Covariates modeling physiological noise sources were included as nuisance regressors during denoising steps (see below).

Denoising was performed by regressing out the confounding effects of the following nuisance regressors: six head motion parameters (translations and rotations used during realignment) and their six first-order temporal derivatives, five principal components modeling physiological noise from within the white matter, five principal components modeling physiological noise from within the CSF, and single volume “scrubbing” regressors modeling outlier volumes that exceeded based on movement and global brain signal thresholds. Following denoising, resting-state data were band-pass filtered between 0.008–0.09 Hz ^27,55^.

A BDC-7 resting-state scan session from one subject (in the iAG group) was excluded from analyses due to excessive movement throughout the run.

#### First-level Analysis

##### Seed-to-Voxel Analysis

We then performed hypothesis-based seed-to-voxel connectivity analyses. This approach estimates resting-state functional connectivity (FC) between *a priori* defined regions of interest (ROIs) and all other voxels in the brain. These ROIs are linked to our hypotheses that AG would have a mitigating effect on HDBR-induced connectivity changes previously reported ^24–26^. We used the same series of ROIs used in our previous bed rest studies ^24–26^; these included regions involved in multisensory integration, somatosensation, motor function, vestibular processing, as well as nodes of the default mode network ^22,26,30–32,56,57^. Table 1 lists each ROI and their MNI space coordinates. Connectivity maps for each ROI were created for each subject and time point using the CONN toolbox. This involved computing the average time series within each (unsmoothed) ROI and calculating the Pearson’s correlation coefficient with data from every other (smoothed) voxel in the brain. Correlation coefficients were Fisher Z-transformed to improve their normality. This procedure was performed for each subject and session.

**Table 1.** Regions of interest (ROIs) used for seed-to-voxel analyses and their MNI space coordinates. L, left; R, right.

| Regions of Interest (ROIs) | MNI Coordinates |  |  |
| --- | --- | --- | --- |
|  | x | y | z |
| R premotor cortex (PMC) | 26 | -10 | 55 |
| R middle frontal gyrus (MFG) | 30 | -1 | 43 |
| L primary motor cortex (M1) | -38 | -26 | 50 |
| L putamen | -28 | 2 | 4 |
| L insular cortex | -41 | 12 | -9 |
| R vestibular cortex | 42 | -24 | 18 |
| L frontal eye field (FEF) | -27 | -9 | 64 |
| L posterior cingulate cortex (PCC) | -12 | -47 | 32 |
| R posterior parietal cortex (PPC) | 28 | -72 | 40 |
| R superior posterior fissure | 24 | -81 | -36 |
| L primary visual cortex (V1) | -10 | -70 | 10 |
| R cerebellar lobule V | 16 | -52 | -22 |
| R cerebellar lobule VIII | 20 | -57 | -53 |

##### Voxel-to-Voxel Analysis

We also used a voxel-to-voxel analysis approach, enabling an exploratory, hypothesis-free assessment of functional connectivity changes with HDBR and whether / how they differ with and without daily AG. This approach calculates an intrinsic connectivity contrast (ICC) reflecting the strength of connectivity between a voxel and all other voxels in the brain ^58^. This involved calculating the root mean square of the Pearson’s correlation coefficients between all voxel time series, then performing a Fisher’s r-to-z transformation. This procedure was performed each resting-state run, resulting in ICC maps reflecting the strength (i.e., magnitude) of connectivity across the brain.

##### Group-level Analysis

###### Longitudinal Changes in Functional Connectivity

We implemented statistical contrasts at the group level to determine the effects of HDBR and whether they were mitigated by daily AG. In our model, our between-subjects factor was group (i.e., cAG, iAG, CTRL) and our within-subjects factor was session (i.e., 5 time points). The same model was used for both seed-to-voxel and voxel-to-voxel analyses. As in our previous bed rest research ^24–26^, our longitudinal model tested for brain networks where functional connectivity was consistent across the first two pre-bed rest time points (i.e., BDC-13 and BDC-7), steadily increased (or decreased) throughout HDBR (i.e., HDT29, HDT58), and then reversed direction, showing recovery towards baseline values on the 10th day after the end of bed rest (i.e., R+10). We evaluated between group effects using an analysis of covariance (ANCOVA) model testing for differences between any of our 3 groups, with subject age and sex used as nuisance covariates. We set our uncorrected voxel threshold at *p* < 0.001 and cluster-level threshold at p < 0.05 corrected for multiple comparisons according to the false discovery rate method. Statistically reliable results were followed up with post hoc t-tests.

###### Brain-Behavior Associations

We performed group-level analyses to test for brain-behavior associations (i.e., correlations between FC changes and performance changes across subjects). Specifically, we investigated whether the 3 groups exhibited differential associations between post-HDBR FC changes and changes in FMT, SOT-5, or SOT-5M scores. Our within-subject contrast modeled FC changes from BDC-7 to HDT58. Our between-subjects regressors of interest modeled the subject-wise performance change per group. Separate models were performed for post-HBDR changes in FMT, SOT-5, and SOT-5M scores. All models included subject age, sex, and group mean were included as covariates of no interest. These analyses allowed us to examine those brain areas in which the association between FC change and mobility or balance changes differed between groups. Results were thresholded as above, and statistically reliable results were followed up with post hoc t-tests.

## RESULTS

### Longitudinal Changes in Functional Connectivity

We first examined if the 3 groups exhibited differential patterns of FC changes from before, during, to after the HDBR intervention.

Our seed-to-voxel analyses revealed a group difference in longitudinal FC changes between our ROI right posterior parietal cortex (PPC) and bilateral clusters centered in the postcentral gyri. Follow-up t-tests revealed that the statistically reliable group difference was between the cAG and CTRL groups (Figure 2). Throughout the HDBR phase, the CTRL group showed increased FC between PPC and bilateral somatosensory cortices, which then decreased to baseline values by 10 days following bed rest. The cAG group showed the opposite pattern, with FC between PPC and somatosensory cortices gradually decreasing by the end of the HDBR phase, followed by a reversal in the post-bed rest recovery phase. Cluster details are shown in Table 2.

**Table 2.** Clusters Showing Differential longitudinal FC changes between cAG and CTRL groups. Peak MNI coordinates and T values of clusters within bilateral clusters in the postcentral gyri (primary somatosensory cortices (S1)) which showed different patterns of FC changes throughout the study between the cAG and CTRL groups. Clusters are FDR corrected. FC, functional connectivity; ROI, region of interest; R, right; L, left.

| ROI | Target cluster | MNI coordinates |  |  | T value | Cluster size |
| --- | --- | --- | --- | --- | --- | --- |
|  |  | x | y | z |  |  |
| R posterior parietal cortex (PPC) | R postcentral gyrus | 58 | -8 | 46 | -6.92 | 96 |
|  | L postcentral gyrus | -46 | -24 | 36 | -6.55 | 58 |
|  | R postcentral gyrus | 62 | -10 | 22 | -6.01 | 46 |

**Figure 2.**
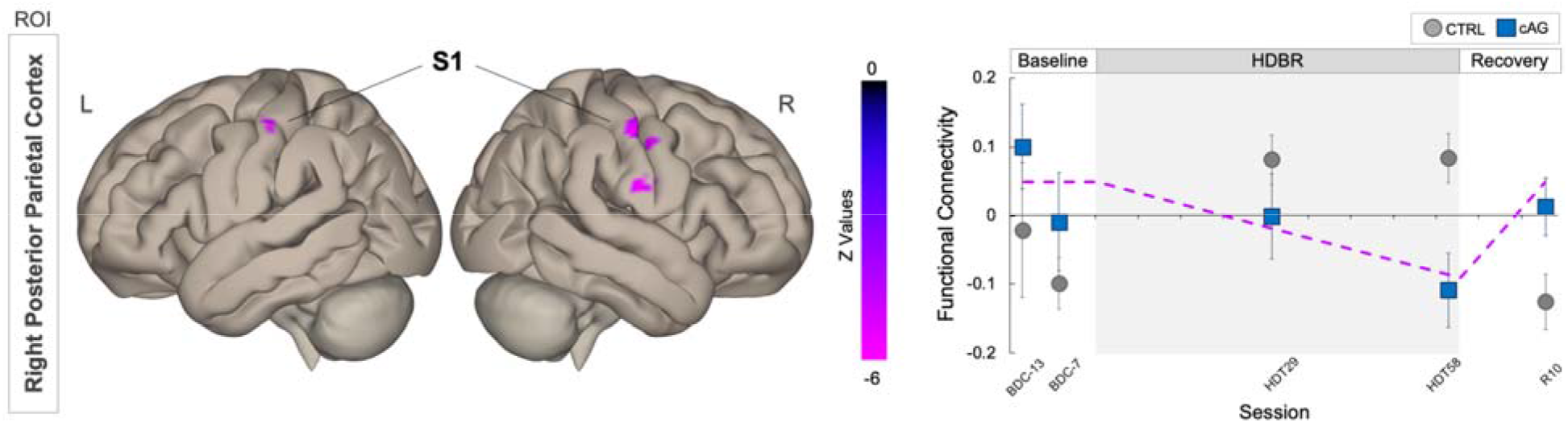
Differential longitudinal FC changes cAG and CTRL groups. The CTRL group (gray circles) showed increased FC between the ROI in right posterior parietal cortex (PPC) and bilateral clusters in primary somatosensory (S1) cortices within postcentral gyri during HDBR followed by a post-bed rest reversal. The cAG subgroup (blue squares) exhibited FC decreases between these regions during HDBR followed by a post-bed rest increase. The dashed line on the graph represents the hypothesized longitudinal model. Error bars represent standard error. L, left; R, right; ROI, region of interest; FC, functional connectivity.

Our hypothesis-free voxel-to-voxel approach did not yield any statistically reliable results.

### Brain-Behavior Associations

We tested whether the 3 groups showed different brain-behavior associations. That is, we evaluated whether there were group differences in the change-change correlation between pre- to post-HDBR FC changes and performance changes in mobility (FMT) or postural control (SOT-5 or SOT-5M).

For FMT total completion time, our seed-to-voxel approach yielded a statistically reliable group difference involving our ROI in left putamen and a cluster in right somatosensory cortex (S1). Follow-up t-tests revealed that the statistically reliable group difference was between the cAG and CTRL groups. This result is shown in Figure 3 and Table 3. Those subjects in the CTRL group who exhibited decreased FC between left putamen and right S1 showed little to no changes in FMT completion time after bed rest while those exhibiting FC increases between these two regions required more time to complete the FMT. Subjects in the cAG group showed the opposite pattern. Within the cAG group, reduced FC between left putamen and right S was associated with greater slowing on the FMT post-HDBR while greater FC was associated with little to no changes in FMT completion time.

**Table 3.** Brain-Behavior Association. Peak MNI coordinates and T values of clusters within right postcentral gyrus that showed differential brain-behavior associations between the cAG and CTRL groups for performance changes on the Functional Mobility Test (FMT).

| ROI | Target cluster | MNI<br>coordinates |  |  | T<br>value | Cluster size |
| --- | --- | --- | --- | --- | --- | --- |
|  |  | *x* | *y* | *z* |  |  |
| L putamen | R postcentral gyrus | 44 | -24 | 40 | -7.22 | 65 |

**Figure 3.**
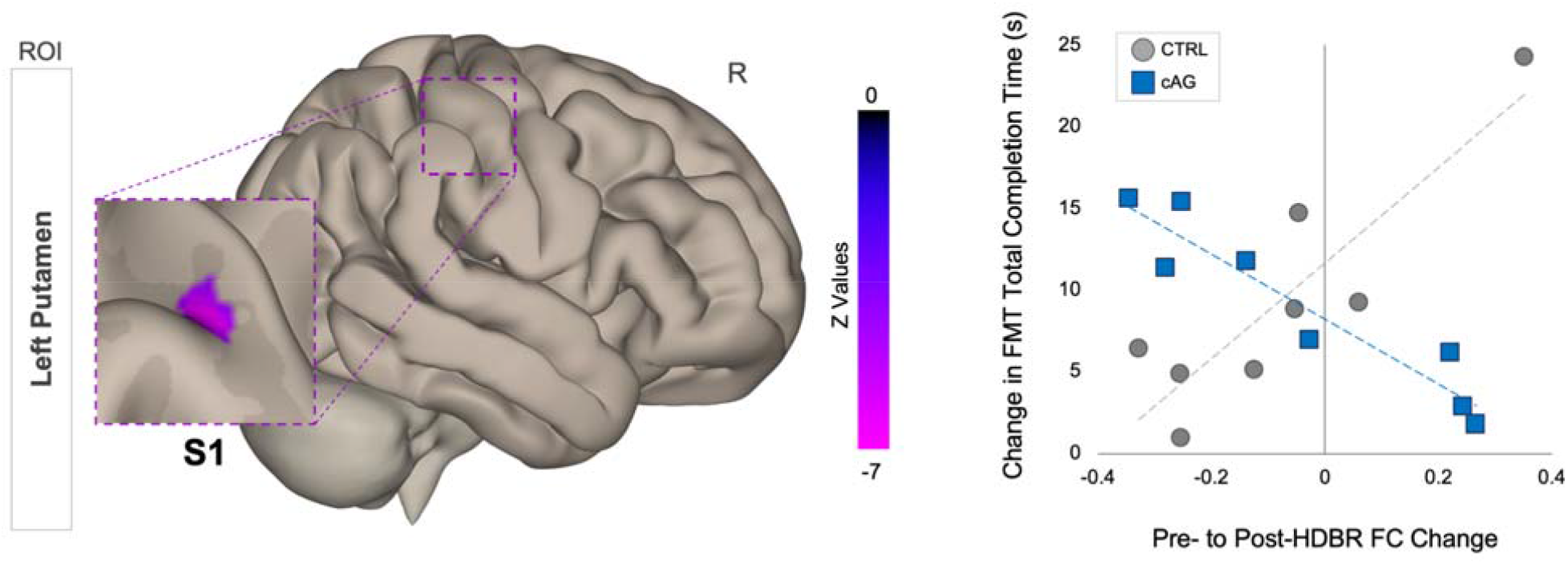
Differential brain-behavior associations between cAG and CTRL groups for the FMT performance. Results of our seed-to-voxel analysis using the ROI in left putamen. Subjects in the CTRL group (gray circles) and cAG group (blue squares) exhibited different patterns of FC change between left putamen and a cluster in primary somatosensory cortex (S1) that were associated with Functional Mobility Test (FMT) completion time changes. On the right, changes in FMT completion time following bed rest are plotted against FC changes from pre-to post-bed rest. R, right; ROI, region of interest; FC, functional connectivity; FMT, Functional Mobility Test; HDBR, head down tilt bed rest.

We observed no statistically reliable group differences in brain-behavior associations relating to post-HDBR changes in postural control measured using either the SOT-5 or SOT-5M. Our hypothesis-free voxel-to-voxel approach did not yield any statistically reliable results.

## DISCUSSION

In the current study, we tested whether and how daily AG exposure administered continuously or intermittently alters resting-state brain functional connectivity during a HDBR spaceflight analog. We additionally tested for group differences in brain-behavior associations involving changes in mobility and postural control from pre-to post-bed rest.

Our hypothesis-driven approach revealed that the cAG and CTRL groups showed differential patterns of FC change throughout the HDBR intervention. The CTRL group showed increased FC between PPC and bilateral S1 during bed rest, which then reversed post-bed rest. The cAG group showed the opposite pattern with gradual FC reductions during HDBR followed by a post-bed rest reversal. Posterior parietal cortex (PPC) is a brain region where multisensory integration occurs, combining inputs from the visual, somatosensory and auditory systems ^59^. Somatosensory cortices receive tactile and proprioceptive inputs and are nodes in densely interconnected networks of brain areas supporting multisensory integration, motor control, and learning ^59–62^. The CTRL group’s FC increase during the bed rest phase likely reflects an increase in gain to compensate for reductions in sensory inputs ^18,63^. During bed rest, subjects experience axial unloading (along the head-to-foot direction), mimicking prolonged periods of reduced sensory inputs from the periphery that occur in microgravity ^64^. Animal models show that peripheral somatosensory deprivation can trigger plasticity within the deprived brain area even in adulthood ^65,66^ and induce fast cortical functional reorganization within surrounding brain areas ^67,68^. Our finding of the opposite pattern in the cAG group suggests that exposure to 30 minutes of continuous axial body loading during centrifugation may have provided peripheral sensory inputs at a sufficient level to mitigate the need to increase somatosensory connectivity (as seen in the control group). This result suggests a differential pattern of somatosensory reweighting throughout HDBR between the cAG and CTRL groups. This is consistent with our hypothesis that restoring afferent inputs with centrifugation would alter HDBR-associated changes in sensorimotor connectivity.

During centrifugation, the magnitude of centrifugal force varies as a function of the distance from the axis of rotation. In this study, subjects in the AG groups experienced ∼2 g_z_ at their feet, providing salient somatosensory inputs at the plantar surface. Despite the magnitude of sensory stimulation in both AG groups, we did not observe differences in the longitudinal pattern of FC changes between the iAG group and the cAG or CTRL groups. It is possible that this null result is attributable to the intermittent AG dosing providing insufficient periods of prolonged somatosensory stimulation. It could also however stem from the small sample size and insufficient power to detect group differences (n=8 per group). Future studies are required to resolve this question.

Somewhat surprisingly, we did not observe group differences in connectivity in brain areas that typically respond to vestibular stimulation, for example retroinsular cortex and parietal opercular area 2 ^11^. It is possible that this may be due to the nature of the HDBR analog and/or the centrifugation procedure used. In HDBR analogs, the subject is reoriented relative to Earth’s gravitational vector, such that the gravitational force in superior-inferior (g_z_; head-to-foot) direction acts in the subject’s anteroposterior (g_x_; front-to-back) direction. During the centrifugation intervention, participants were exposed to hypergravity: centrifugal force applied along the head-to-foot direction (g_z_) in addition to Earth’s gravity acting along the participants’ front-to-back (g_x_) direction. Moreover, due to the centrifugation procedure, the gravitational vector acting on the head (and vestibular apparatus) would still be biased towards g_x_. Positioned closer to the axis of rotation, the head and vestibular apparatus was exposed to approximately 0.3 g along the longitudinal (z) axis ^69^, which may not have provided sufficient vestibular activation. As these are unavoidable shortcomings of the HDBR model and centrifugation, data from spaceflight is required to fully address whether or how vestibular stimulation during centrifugation alters FC in microgravity.

We further found that the cAG and CTRL groups exhibited distinct brain-behavior associations with respect to mobility changes following bed rest. For the CTRL group, reductions in FC between left putamen and right S1 after HDBR were associated with smaller decrements in mobility post-bed rest. For the cAG group, FC increases between these brain areas after HDBR were associated with smaller decrements in mobility. The observed brain-behavior relationship suggests that, when somatosensory stimulation is made available, FC increases between the putamen and S1 during bed rest are compensatory in nature, reducing post-HDBR mobility declines. The somatosensory cortex and the putamen are connected via thalamocortical striatal pathways ^70,71^, and conditions that impact striatal function, such as Parkinson’s disease, affect somatosensory function ^72^. Thus, there is an anatomical basis for these effects.

A recent study of 11 cosmonauts reported task-based FC changes among sensorimotor, visual, vestibular and somatosensory/proprioceptive brain areas during foot stimulation following long-duration spaceflight, suggestive of post-flight somatosensory and vestibular reweighting ^13^. Without accompanying measures of performance change post-flight ^13^, the nature of these functional brain changes (i.e., adaptive or dysfunctional) is unclear ^18^. In a previous 70-day bed rest study, our group found that a similar region of S1 (among other brain areas) was activated during a foot tapping task, and greater S1 activation during a foot tapping task at the end of bed rest was associated with smaller decrements in functional mobility performance ^73^. This finding is suggestive of compensatory sensorimotor reweighting during HDBR, with greater sensorimotor resources required for both foot movement and mobility after bed rest.

## Limitations

A few limitations of this study should be noted. Group sample sizes (n=8) were small, reducing the sensitivity of our analyses. We also had fewer fMRI sessions throughout the study compared to previous bed rest campaigns ^25,26^, with only one session during the recovery phase occurring 10 days after subjects had exited bed rest. This reduced our ability to detect faster FC changes that might have occurred shortly after exiting bed rest; this leaves the question unresolved as to whether there are differential recovery rates for those that did versus did not receive AG during HDBR. Additionally, as noted above, subjects received less than 1 g_z_ at their head during centrifugation, which may have limited vestibular stimulation during AG. Indeed, we had expected to see effects of AG on vestibular cortical connectivity patterns, but we did not.

## Conclusions

Here we have shown that 30 minutes of continuous AG exposure during HDBR alters FC changes between posterior parietal cortex and primary somatosensory (S1) cortices. Additionally, the cAG and CTRL groups showed different brain-behavior correlations between the putamen and S1 associated with post-bed rest changes in mobility, suggestive of sensory reweighting. The S1 clusters are located within regions we have previously shown to be activated during foot movement and that exhibit mobility-related changes following bed rest ^73^. This suggests that the observed FC changes in the cAG group may have been due to foot somatosensory stimulation. The feet were exposed to approximately 2 g_z_ during AG, providing salient somatosensory stimulation during HDBR. Resting-state FC changes during the HDBR phase differed only between the cAG and CTRL groups. This may point to differences in the effectiveness of centrifugation delivery (intermittent vs. continuous) on stimulating the brain’s somatosensory system and inducing sensory reweighting. However, additional work is needed to determine if this reflects our small sample size and limited statistical power.

